# Color discrimination in action: High-throughput measurement in immersive VR

**DOI:** 10.64898/2026.02.13.705758

**Authors:** Giulia Agosti, Jacob Hadnett-Hunter, Karl R. Gegenfurtner

**Affiliations:** Abteilung Allgemeine Psychologie and Center for Mind, Brain & Behavior Justus-Liebig-Universität Gießen, Gießen, Germany

## Abstract

Precise measurement of color discrimination across color space is limited by the time and effort required to collect large psychophysical datasets. We investigated whether immersive virtual reality (VR) can support high-throughput measurement of color discrimination without compromising data quality. A standard 4-alternative forced-choice odd-one-out task was embedded in an interactive VR environment inspired by the rhythm game *BeatSaber*, in which participants indicated the location of a chromatic target by slicing approaching cubes.

Stimuli were presented on a color-calibrated VR headset, and chromaticities were specified in DKL space. Discrimination thresholds were measured for hue and chroma shifts around two reference colors. Participants sustained response rates of approximately one trial per second while maintaining stable performance.

Thresholds replicated established asymmetries in color discrimination: hue thresholds were lower than chroma thresholds, and the hue–chroma ratio differed between color quadrants. A control experiment comparing VR-based slicing responses with matched keyboard responses revealed comparable psychometric fits and threshold estimates, indicating that motor engagement did not degrade measurement precision. Questionnaire measures further showed significantly higher intrinsic motivation, enjoyment, and stimulation in the immersive condition relative to a classic static psychophysical task, without increases in reported pressure or discomfort.

These results demonstrate that calibrated immersive VR can yield reliable color discrimination measurements at substantially increased throughput, providing a scalable approach for mapping the metric structure of color space.

## Introduction

Color discrimination is a fundamental aspect of human visual perception. Our ability to differentiate between subtly different colors is crucial for tasks ranging from object recognition to aesthetic appreciation. Color discrimination has been studied for more than a century (for a historical review, see Mollon & Danilova, 2025), using both theoretical modelling (Schrödinger, 1920; Stiles, 1959; Von Helmholtz, 1867), and empirical approaches (MacAdam, 1942; Wyszecki & Fielder, 1971; Wright, 1941; Krauskopf & Gegenfurtner, 1992; Rinner & Gegenfurtner, 2000; M. V. Danilova & Mollon, 2012, 2014). These studies have provided important insights into the mechanisms underlying color vision and have established a rich tradition of psychophysical measurement.

Color discrimination thresholds have traditionally been measured using color-matching procedures (Brown & MacAdam, 1949; MacAdam, 1942) or odd-one-out paradigms (Krauskopf & Gegenfurtner, 1992; Rinner & Gegenfurtner, 2000; Hansen et al., 2008; Hedjar et al., 2025a, 2025b), from which discrimination contours can be derived. Such measurements also form an important empirical foundation for color-difference formulas such as CIEDE2000 (Fairchild, 2013; Gao et al., 2023; Luo et al., 2001, 2023; Sharma et al., 2005). Despite the success of these approaches, the available discrimination data remain relatively sparse compared with the size and complexity of color space, largely because threshold measurements are time-consuming and require many repeated observations. Here, we explore whether immersive virtual reality can increase the efficiency of psychophysical data collection while preserving the quality of color-discrimination measurements.

The difficulty of measuring the full metric of color space is best illustrated by the enormous effort David MacAdam (MacAdam, 1942) undertook to determine the seminal discrimination ellipses that bear his name. These remain among the most complete direct measurements available, even though thresholds were measured only in a single plane of color space. Later, MacAdam and Brown (Brown & MacAdam, 1949) and Wyszecki and Fielder (Wyszecki & Fielder, 1971) extended the work to three-dimensional discrimination ellipsoids, still limited to the same chromaticity centres. The study of Brown and MacAdam (Brown & MacAdam, 1949) required about 48,000 individual settings over 18 months of testing by a single observer. Clearly, extending this approach to the entire three-dimensional color space is not feasible. The space is vast, sensitivity varies with direction and location, and the number of required measurements grows exponentially with dimensionality; a classic example of the “curse of dimensionality” (Bellman, 1957). As a result, no comprehensive dataset exists that maps discrimination thresholds across the full color space.

What can be done then? One approach is to increase data throughput by using paradigms that allow faster sequences of judgements and are less tiring for the observer. Examples include continuous psychophysics ( Barnett et al., 2025; Bonnen et al., 2015, 2017; Huk et al., 2018; Knöll et al., 2018; Straub & Rothkopf, 2022), in which observers continuously track changing stimuli and perceptual performance is quantified from the resulting response trajectory, as well as immersive and game-based paradigms such as the one explored here. A different approach seeks to reduce the number of required measurements through interpolation or global fitting. Hong et al., for example, used advanced fitting methods to reconstruct a complete metric field from a limited set of measurements, albeit within a single plane of color space (Hong et al., 2025). Here, we pursue the first strategy by embedding a standard color-discrimination task within an interactive VR game that participants can perform for extended periods of time. In the game, as in the real world, perceptual judgements are embedded within action and interaction. In principle, increased-throughput and model-based approaches are complementary and could ultimately be combined.

Our approach also addresses broader questions about the generalizability of laboratory psychophysics. While highly controlled experiments have provided foundational insights into perception, it remains important to assess whether established psychophysical measurements are preserved in more naturalistic settings (Kingstone et al., 2008; Osborne-Crowley, 2020; Shamay-Tsoory & Mendelsohn, 2019). In everyday vision, perceptual judgments are embedded within action and interaction with the environment. Immersive virtual reality provides an opportunity to study perception under such conditions while retaining precise control over stimulus presentation.

We therefore measured color discrimination thresholds in an immersive, action-based task first introduced by Aizenman et al. <u>(2026)</u> to study the stability of color categories. Although absolute thresholds are expected to depend on experimental details such as adaptation state and stimulus timing, several qualitative features of color discrimination are known to be robust. In particular, discrimination contours are anisotropic, with greater sensitivity to hue than to chroma changes, and this anisotropy varies systematically across color space. MacAdam’s <u>(1942)</u> measurements revealed elliptical discrimination contours in chromaticity space. Later work showed that thresholds are typically lower for hue than for chroma changes, a phenomenon Judd (1969, 1970) termed the “super-importance of hue,” and that the strength of this asymmetry varies across color space (M. V. Danilova & Mollon, 2012, 2014, 2020; Hedjar et al., 2025a, 2025b). The central question here was whether these canonical patterns would be preserved under immersive, motor-engaged conditions.

## Main Experiment

To address this question, we embedded a standard 4-AFC odd-one-out color discrimination paradigm within an immersive virtual reality (VR) environment inspired by the rhythm game *BeatSaber*. Here, we used the “Saberception” toolbox (Hadnett-Hunter et al., 2026) to quantify color discrimination thresholds. On each trial, four color patches were presented on the front face of an approaching cube. Three patches shared a reference color, and one was a target differing in either hue or chroma. Participants indicated the location of the odd-one-out target by slicing the cube in the corresponding direction. By systematically varying the chromatic difference between reference and target using a method-of-constant-stimuli procedure, we estimated hue and chroma discrimination thresholds.

### Methods

#### A. Participants

A total of 26 participants (15 females, average age = 29.8, range 23-38 years) took part in the first experiment. Two participants had to be excluded because they did not follow the task instructions correctly. All participants had normal or corrected-to-normal vision and normal color vision, as tested by the 24-plate edition of Ishihara plates (Clark, 1924). Before the experiment, participants were informed about the procedures and signed written consent forms. This study was approved by the ethics commission of the Department of Psychology at Justus-Liebig-Universität Giessen (LEK 2020-0015).

#### B. Apparatus

We developed our ‘BeatSaber’ clone using Unity (version 6000.0.22f1). The game was built for Android to run locally on Meta Quest headsets and made use of the Meta XR Core, Interaction and Haptics SDKs (version 76.0). For our experiment, we presented the game on a Meta Quest 3 headset (90 Hz refresh rate, LCD display, 8-bit color depth, resolution of 2064 x 2208 pixels per eye). The display was color calibrated by a CS-2000A spectroradiometer (Konica Minolta, Chiyoda, Tokyo, Japan), with calibration stimulus control being implemented in Unity and MATLAB, following prior work by Díaz Barrancas et al. (Díaz-Barrancas et al., 2024; Díaz–Barrancas et al., 2023). The red, green, and blue primaries of the display had CIE *x*, *y*, *Y* coordinates of [0.649, 0.327, 98.337], [0.299, 0.606, 346.140], and [0.151,0.058, 35.606], respectively.

Color discrimination thresholds were measured across eight conditions defined by two reference colors (purple-ish and orange-ish), two stimulus dimensions (hue and chroma), and two shift directions for each dimension: toward or away from the grey adaptation point for chroma, and clockwise or counter-clockwise angular shifts for hue (see Figure 1).

**Fig 1.**
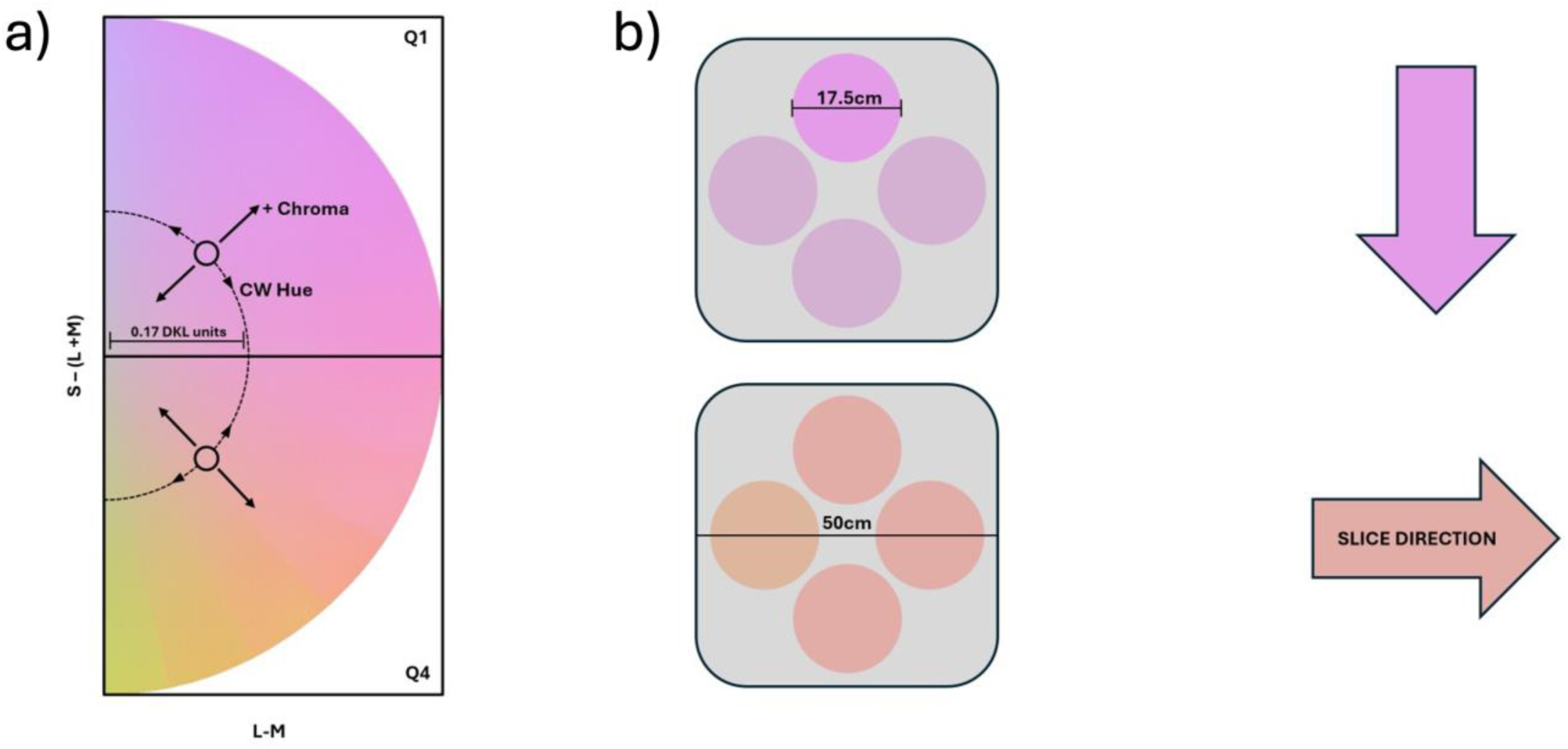
(a) Reference colors (black circles) and the radial (chroma) and tangential (hue) directions along which discrimination was measured in the DKL isoluminant plane. (b) Schematic of the stimulus configuration on the front face of the approaching cube. Three patches were presented in the reference color and one patch was an odd-one-out target. Participants indicated the target location by slicing the cube from the side containing the target patch.

We specified chromaticities in Derrington–Krauskopf–Lennie (DKL) (Derrington et al., 1984) color space and we used conventions detailed in Hansen and Gegenfurtner (2013) to define the DKL axes and to compute the transformation from DKL to device RGB values.

#### C. Stimuli and Procedure

The two reference colors lay on radial lines in the DKL isoluminant plane, one at 45° (Quadrant 1, purplish) and the other at 315° (Quadrant 4, orangish). The reference colors lay 0.17 DKL units from the origin. This eccentricity was chosen to be clearly suprathreshold while remaining sufficiently far from the gamut boundaries to allow comparison stimuli to be generated in all directions without clipping. For each discrimination dimension (hue, chroma) and direction, we computed 10 shifts between 0 and 0.07 DKL units from the reference point. For chroma, shifts were defined as radial displacements either toward or away from the neutral point. For hue, shifts were defined as tangential displacements corresponding to clockwise or counter-clockwise movement around the isoluminant plane (Figure 1a). On each trial, four color patches were displayed on the front face of an approaching cube. Three patches shared the reference color and one was an odd-one-out target. Participants indicated the location of the target by slicing the cube from the side containing the target patch. Thresholds were estimated using the method of constant stimuli.

Figure 2 shows the progression of a single cube during a trial (see also Movie 1). Each trial consisted of participants slicing a cube moving towards them that contained four color patches on the front face. Three patches were the reference color and one odd-one-out was a shifted color. The cube used an unlit material set to the isoluminant mid-grey color. Cube faces were approximately 50cm in width and height, and each patch was 17.5cm in diameter. During the stimulus presentation range, this meant that patches subtended approximately 1 – 2 degrees of visual angle. Each cube was generated at a fixed distance of 40m from the participant’s viewpoint and moved along a predefined trajectory that ended with a final “bounce” towards the player. We defined a trial as the 1-second interval from the onset of the bounce to the moment it was slashed by the participant. In the first 500ms of this interval, the block displayed the stimuli. After this, mid bounce, the cube’s face turned blank. The cube continued its trajectory towards the participant for another 500ms during which it could be sliced. Participants received auditory feedback upon slicing the cube.

**Fig 2.**
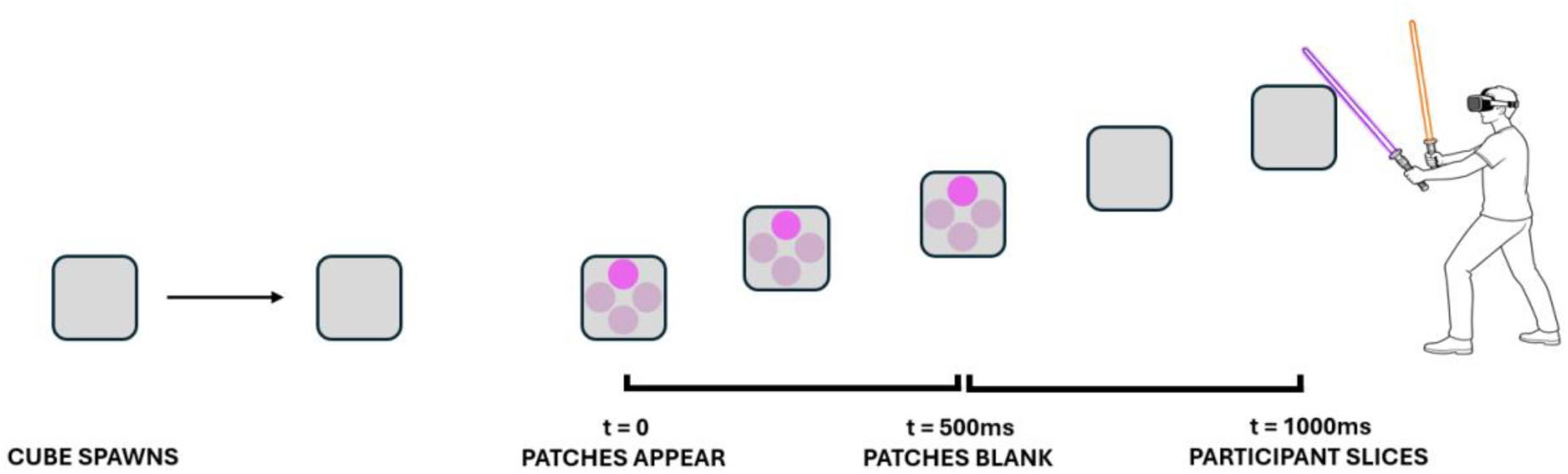
Example progression of a single cube during a trial.

**Movie 1.**
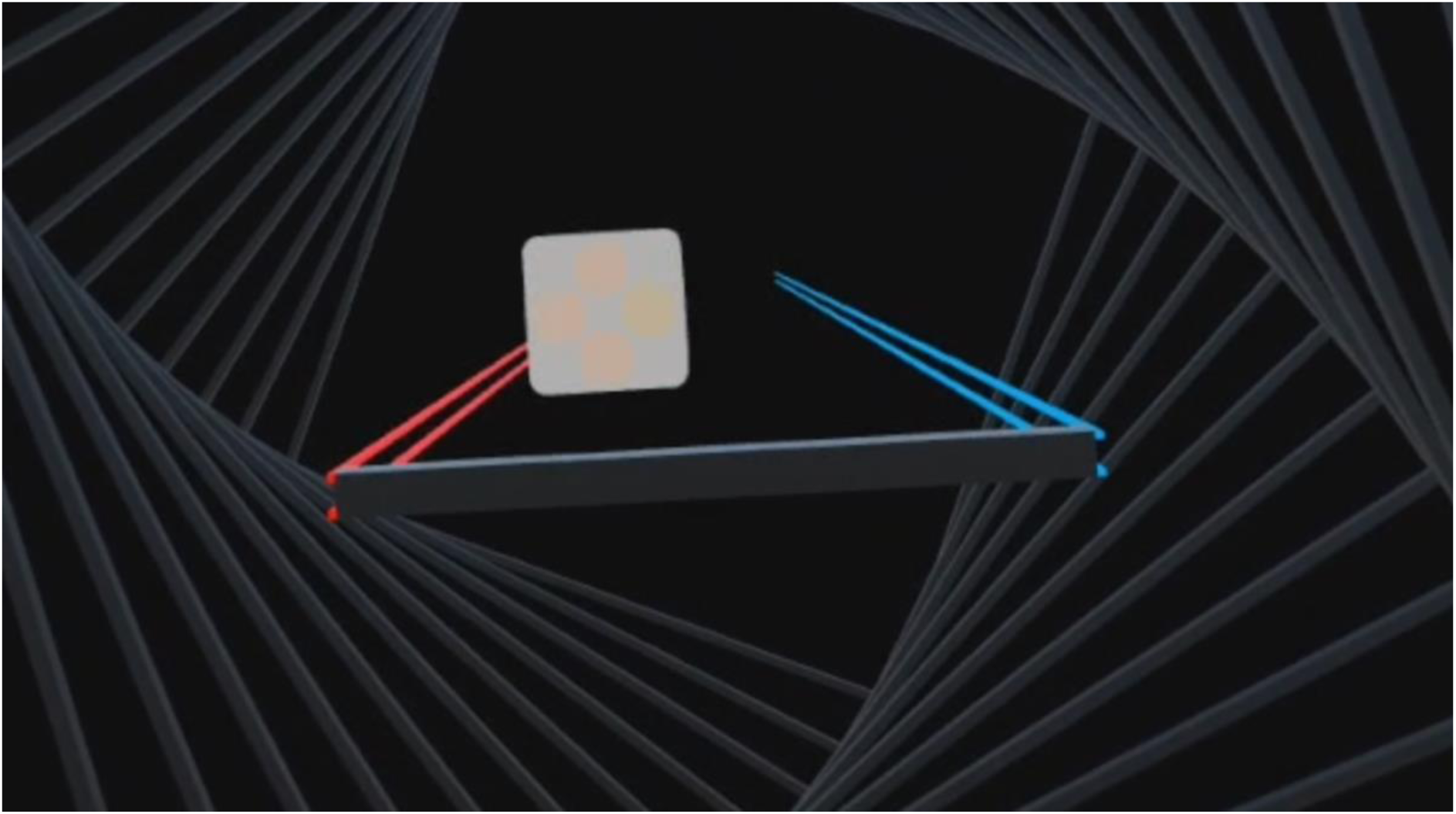
A participant playing the BeatSaber game. See uploaded movie file.

Blocks of trials were organized into 2.5-minute soundtracks. Participants initiated each block autonomously via a start menu. Figure 3 shows an example stream of trials within a single soundtrack. Each soundtrack contained 100 trials split between two conditions in separate left and right cube streams. Each stream contained 50 cubes, 5 repetitions of each of the 10 target patch intensities. The soundtracks were all 160 beats per minute, with cubes synchronized to appear on every 4th beat, resulting in 40 trials per minute.

**Fig 3.**
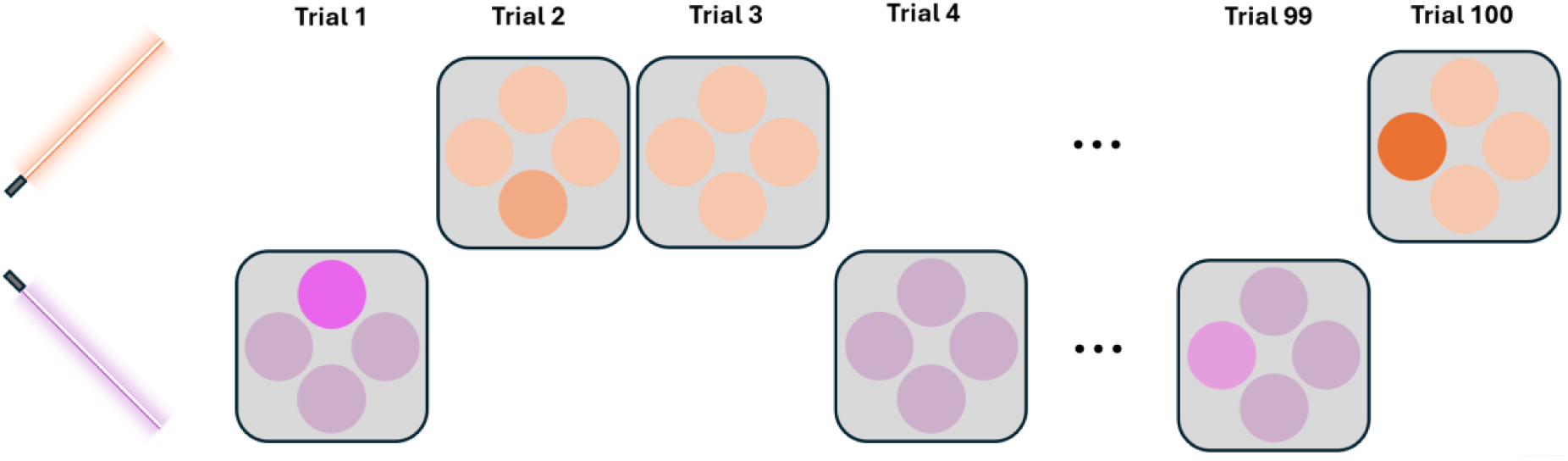
Example stream of cubes (trials) in a block. Each block contained 100 trials. Color conditions were split into left and right streams, 50 on either side.

Prior to the main experiment, participants completed a short training session to gain familiarity with the task. The training session required participants to complete soundtracks in which the target patch was highly different from the reference. They played until they achieved a hit rate of 97% or above. For the main experiment, we separated the 8 conditions into 4 soundtracks, each containing an orangish and purplish stream and with target patch shifts being either in hue clockwise, hue counter-clockwise, chroma positive, or chroma negative. Participants completed each soundtrack 3 times, resulting in 15 repetitions per stimulus intensity level for all 8 conditions.

Finally, to ensure that the ‘BeatSaber’ methodology did not produce unwanted discomfort or sickness for participants, they completed the Simulator Sickness Questionnaire (Kennedy et al., 1993) at the end of the experiment.

### Results and Discussion

We fitted psychometric functions to individual participants using the psignifit toolbox for MATLAB (Schütt et al., 2016; Wichmann & Hill, 2001a). The lower asymptote was fixed at 25% correct (chance performance in a 4AFC task). Discrimination thresholds were defined as the stimulus difference corresponding to 62.5% correct performance. Figure 4 shows representative psychometric function fits. Performance increased smoothly with stimulus difference, indicating that the 8-bit display resolution did not limit the measurements.

**Fig 4.**
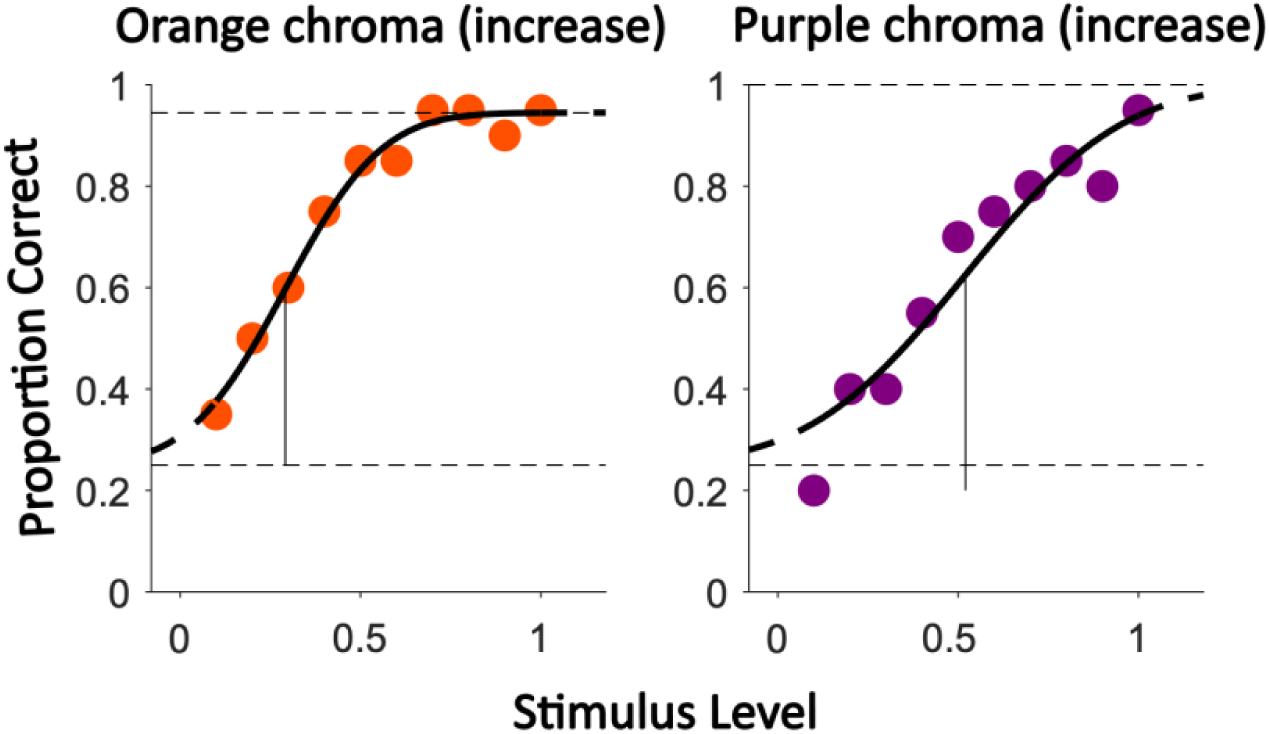
Exemplary individual psychometric curve fits for orange (left) and purple (right), for a single direction. The x-axis represents stimulus intensity in discrete steps along the DKL color space, measured relative to the reference color at 0. The y-axis represents the proportion of correct target selections. The colored circles represent data for each condition, and solid black curves show the fitted psychometric functions. The vertical grey lines indicate the estimated discrimination thresholds (62.5% correct). Dashed horizontal lines indicate chance performance (25%) and the estimated upper asymptote.

Figure 5 shows discrimination thresholds for the purple and orange reference colors in the DKL isoluminant plane. Thresholds were estimated separately for hue and chroma and then averaged across both directions within each dimension (i.e., increasing and decreasing chroma; clockwise and counter-clockwise hue shifts).

**Fig 5.**
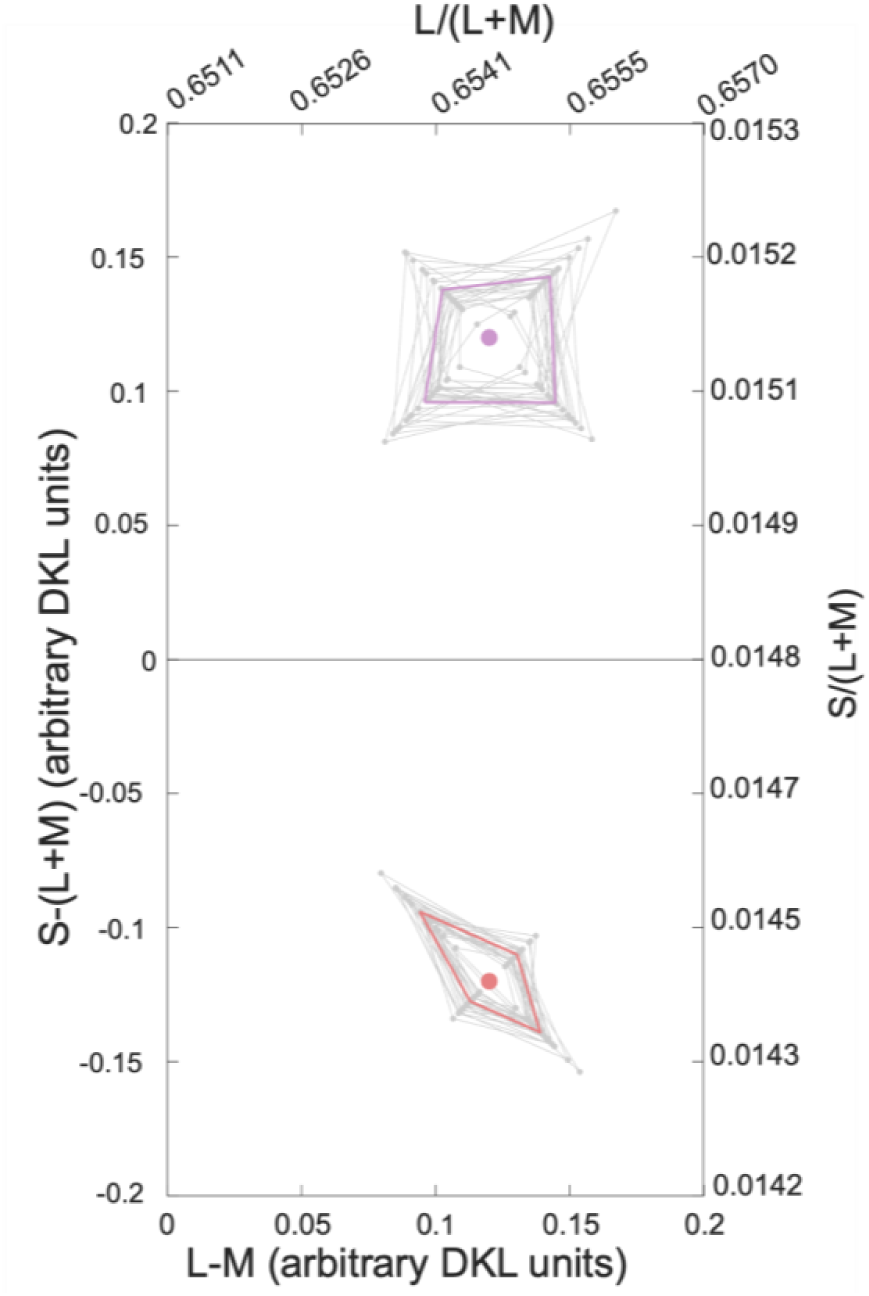
Individual discrimination thresholds for all participants shown in the DKL isoluminant plane. The two reference colors are indicated by the colored dots. Colored contours show mean discrimination thresholds and grey contours the thresholds of individual participants. The upper and right axes indicate the corresponding coordinates in MacLeod–Boynton space (MacLeod & Boynton, 1979). DKL coordinates are shown in the units defined by the transformation described in Hedjar et al. (Hedjar et al., 2025b).

To assess the effects of reference color and discrimination dimension, we performed a 2 × 2 repeated-measures ANOVA with the within-subject factors Quadrant (Purple, Orange) and Color Dimension (Hue, Chroma). We tested for main effects of both factors as well as their interaction. Significant effects were followed by pairwise comparisons to characterize the observed differences.

Figure 6 summarizes discrimination thresholds for hue and chroma around the orange and purple reference colors. Consistent with previous work (Hedjar et al., 2025a, 2025b; MacAdam, 1942), discrimination was anisotropic, with larger thresholds along the chroma than the hue direction. A 2 × 2 repeated-measures ANOVA revealed significant main effects of Color Dimension (F(1,23) = 44.65, p < 0.001) and Quadrant (F(1,23) = 65.24, p < 0.001), as well as a significant interaction between the two factors (F(1,23) = 38.14, p < 0.001). The interaction reflects the fact that the hue–chroma difference depended on reference color. Post hoc comparisons showed that chroma thresholds were significantly larger than hue thresholds in the orange quadrant (*p* < 0.001), whereas the corresponding difference in the purple quadrant did not reach significance (*p* = 0.098). Comparing quadrants within each discrimination dimension revealed no significant difference for chroma thresholds (*p* = 0.63), but significantly lower hue thresholds in the orange than in the purple quadrant (*p* < 0.001).

**Fig 6.**
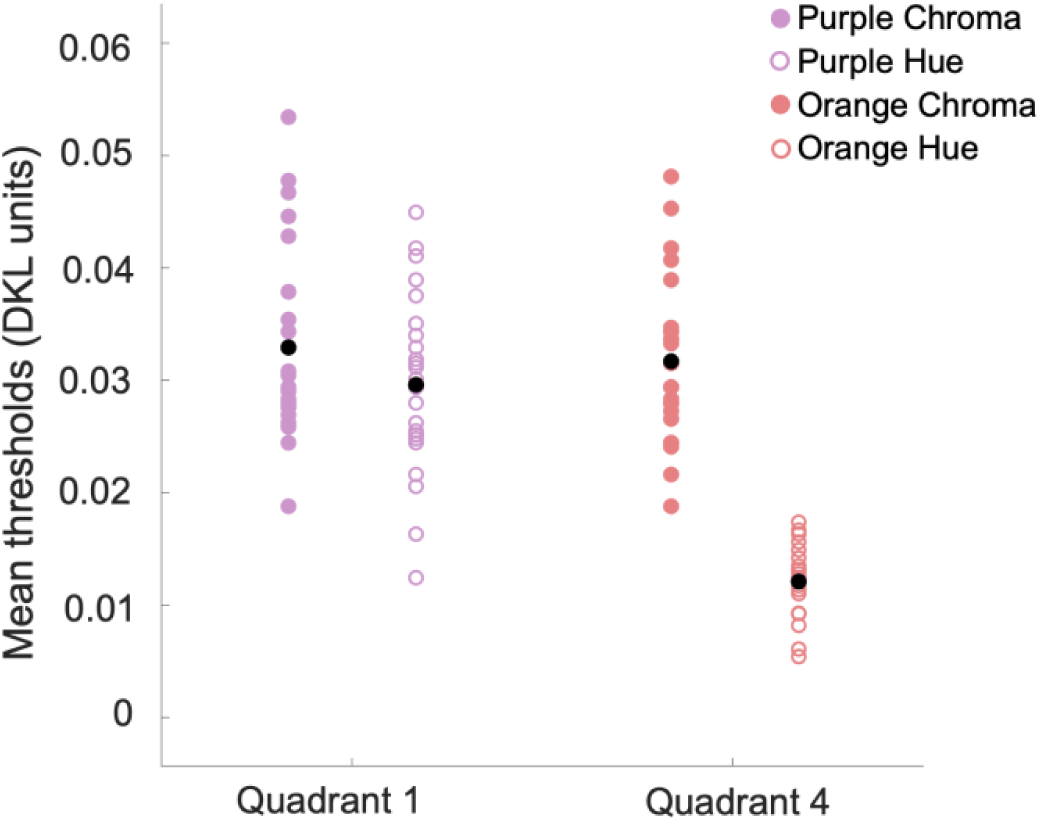
Mean and individual discrimination thresholds expressed as Euclidean distance from the reference colors in the DKL isoluminant plane. Results are shown separately for hue and chroma in the purple (Quadrant 1) and orange (Quadrant 4) regions. Each participant is represented by a colored symbol (filled: chroma; open: hue), and group means are indicated by black symbols.

Our study replicated well-established asymmetries in color discrimination. In the DKL isoluminant plane, discrimination contours were elongated predominantly along radial (constant-hue) directions. Moreover, this elongation varied systematically across color space and was markedly stronger for the orange than for the purple reference color. While the numerical magnitude of hue–chroma threshold ratios depends on the color-space representation, DKL provides a physiologically motivated coordinate system whose axes approximate the cardinal post-receptoral chromatic mechanisms. Within this framework, our results are consistent with Judd’s notion of the “super-importance of hue” (Judd, 1969, 1970; Danilova & Mollon, 2016) and demonstrate that this classical asymmetry remains robust under immersive, action-based viewing conditions.

To assess whether VR-related discomfort might have influenced participant performance, we administered the Simulator Sickness Questionnaire (SSQ) at the end of the experimental session. The SSQ yields a composite score ranging from 0 to 235.6, with higher values indicating greater simulator sickness (Kennedy et al., 1993). The mean SSQ score was 21.14 ± 19.99. In comparison, the meta-analysis by Saredakis et al. (Saredakis et al., 2020) on visually induced motion sickness reported a higher average of 27.4 ± 3.1 for experiments of similar length. In addition, no participant reported discomfort or simulator sickness during the experiment. These results suggest that the head-mounted display (HMD) and immersive VR experimental design did not elicit significant visual discomfort during the study.

A key contribution of our study is the integration of psychophysical measurement within an engaging, game-based task. Gamified paradigms have been shown to enhance participant motivation, enjoyment, and sustained attention during perceptual and cognitive tasks (Hamari et al., 2014; Lumsden et al., 2016; Mekler et al., 2017). For psychophysics, such factors are not merely experiential but methodological: reduced boredom and attentional lapses may stabilize performance across extended testing sessions and improve effective data yield. In the present task, minimal training was required, and participants were able to sustain a high trial rate while maintaining stable psychometric performance throughout a session. Lapse rates were also very low (< 5%) throughout the experiments.

While these findings demonstrate that color discrimination can be measured effectively in an immersive, game-based environment, it remains essential to establish that the paradigm does not compromise psychometric quality or introduce systematic bias. To address this question, we conducted two control experiments that compared the reliability, efficiency, and subjective experience of the BeatSaber paradigm with more traditional response methods.

## Control experiments

We designed two additional control conditions to evaluate both the psychometric properties and the subjective experience of the BeatSaber paradigm. The purpose of the control experiments was not to reproduce the full discrimination dataset, but to evaluate measurement reliability, throughput, and participant experience across paradigms. The first control condition (Control Key-Press) differed only in the motor response required. The second control condition (Control Classic) more closely resembled a conventional psychophysical color discrimination experiment. It allowed us to assess the extent to which the increased enjoyment reported for BeatSaber reflected the broader game environment rather than the response modality alone.

### Methods

10 participants from the main experiment returned to take part in the control experiments. In one session we measured color discrimination thresholds using the ‘BeatSaber’ method, Control Key-Press, and Control Classic. The order of the experimental conditions was counterbalanced across participants, but they had all played BeatSaber earlier in the main experiment.

To reduce testing time, thresholds were measured for a single hue direction and a single chroma direction around each reference color. Specifically, we measured thresholds for decreasing chroma and counter-clockwise hue shifts. For each experimental condition (BeatSaber, Control Key-Press, Control Classic) we collected 600 trials, so that we had the same number of repetitions as in the main experiment.

In the Control Key-Press condition, the visual stimuli, timing, and task structure were identical to BeatSaber, but participants reported the location of the odd-one-out target using keyboard arrow keys rather than arm movements. Because the moving cubes, music, and game-like presentation might themselves influence engagement, the Control Classic condition more closely resembled a conventional psychophysical color-discrimination experiment. Participants responded with arrow keys to static color patches presented centrally on a uniform grey background. No music was played, and the dynamic game elements were removed. The presentation time was matched to the duration of BeatSaber and Control Key-Press trials (500ms). Stimulus size matched the apparent size of the color patches at the moment of response in the BeatSaber condition (approximately 2° visual angle). No music was played for this condition, and responses were given via key press, with only feedback sounds provided.

In BeatSaber, trials ran continuously from beginning to end of each block. In both Control paradigms the experiment paused on each trial until a response was made, a trial structure consistent with standard practice in classical psychophysical experiments. As a result, blocks in the BeatSaber condition had a fixed duration of 150 seconds, whereas block duration in the Control conditions varied depending on participants’ response times. This difference in trial structure allowed us to assess whether continuous stimulus presentation contributes to the throughput advantage of the BeatSaber paradigm. After completing each condition, participants filled out the Intrinsic Motivation Inventory (IMI) (Ryan et al., 1983) and the User Experience Questionnaire (UEQ) (Laugwitz et al., 2008).

From the original IMI, we used a version with selected items from the Interest/Enjoyment and Pressure scale. It consisted of 12 items scored on a Likert scale from 1 (not at all) to 7 (very true).

The UEQ is comprised of 6 scales (Attractiveness, Perspicuity, Novelty, Stimulation, Dependability, Efficiency). Items have the form of a semantic differential, i.e. each item is represented by two terms with opposite meaning and are scored on a seven-stage scale. We adapted the UEQ introduction by replacing the term ‘product’ with ‘experiment’.

### Results and Discussion

We first assessed the quality of the data collected across the three conditions – BeatSaber, Control Key-Press and Control Classic. To evaluate goodness of fit, we computed the deviance between the observed responses and the fitted psychometric function as described by Wichmann & Hill (Wichmann & Hill, 2001a, 2001b). Under the assumption that responses follow binomial variability around the fitted model, the deviance is approximately χ²-distributed with degrees of freedom equal to the number of stimulus levels minus the number of free parameters. We therefore compared each deviance value to the corresponding χ² distribution to obtain a p-value, and fits with p < 0.05 were considered to show significant lack of fit.

Across the 40 psychometric fits obtained in each condition, only a small number showed significant lack of fit (BeatSaber: 4; Control Key-Press: 1; Control Classic: 4). These rates are comparable to those typically observed in psychophysical datasets (Jäkel & Wichmann, 2006) and indicate that the fitted psychometric functions provided an adequate description of observer performance in all three conditions.

Figure 7 compares individual and mean chroma and hue thresholds obtained with the BeatSaber, Control Key-Press, and Control Classic paradigms. The diagonal error bars represent 95% confidence intervals of the mean difference (Schütz & Gegenfurtner, 2025). Thresholds obtained with BeatSaber and Control Key-Press were highly similar, indicating that replacing the slicing response with a keyboard response had little effect on measured color-discrimination thresholds. Although BeatSaber and Control Classic differed in several aspects of stimulus presentation and adaptation state, the overall pattern of thresholds was also broadly comparable across these paradigms.

**Fig 7.**
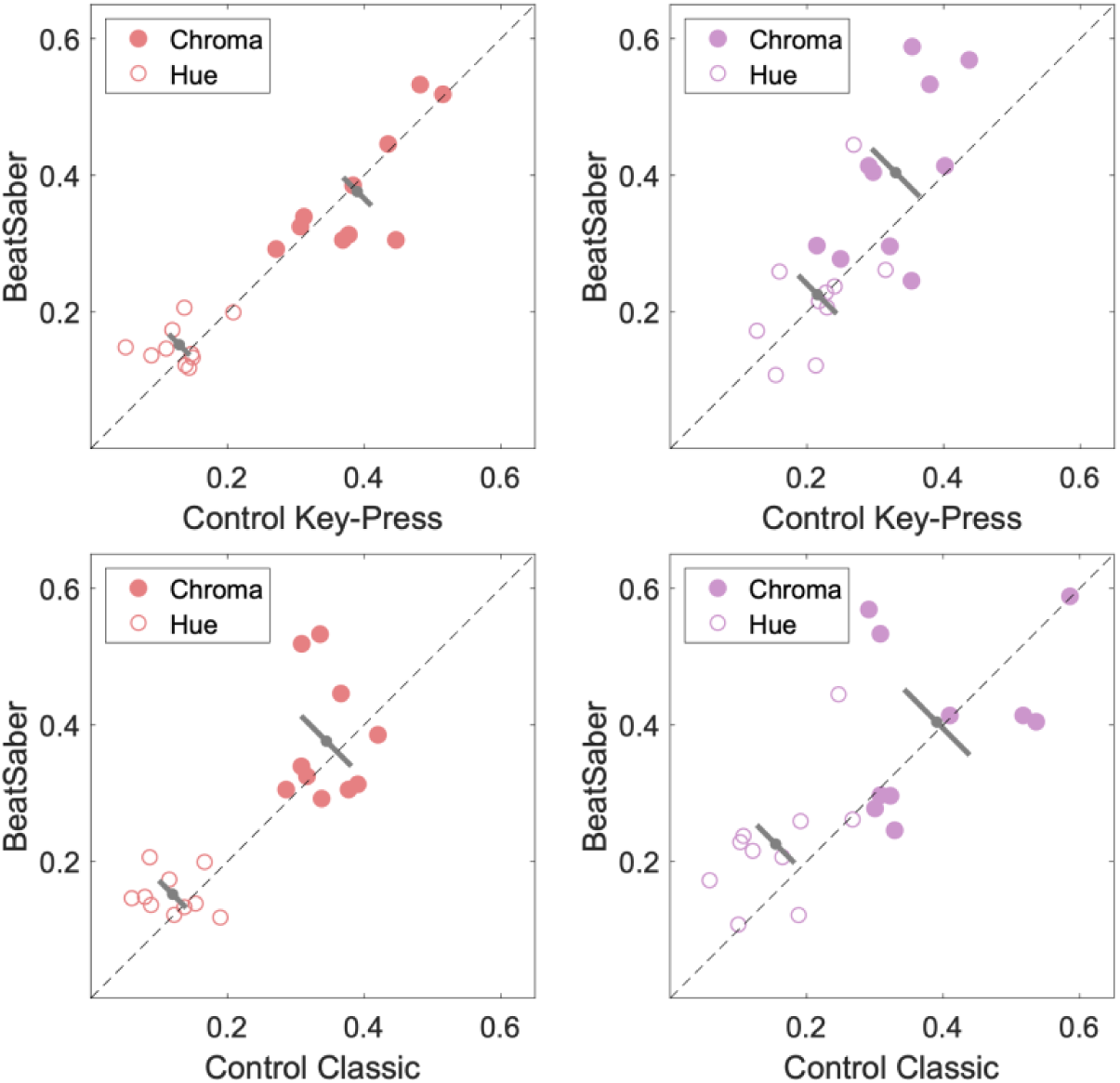
Discrimination thresholds for chroma and hue for the BeatSaber and Control Key-Press (top row) and BeatSaber and Control Classic (bottom row) conditions. Axes show normalized discrimination thresholds, where 0 corresponds to the reference color and 1 to the largest tested stimulus difference. Left panels display thresholds for the purple quadrant and right panels for the orange quadrant. Filled symbols represent chroma thresholds and open symbols hue thresholds for individual participants. Grey diagonal bars indicate the group mean difference and its 95% confidence interval.

Based on timestamps recorded at the beginning and end of each block, we calculated the total time required to complete a block of 100 trials. These measurements included the time needed to select and initiate each block from the experiment menu, which was identical across paradigms.

Figure 8 shows the average completion times. BeatSaber blocks required 158 s on average, compared with 202 s and 206 s for the Control Key-Press and Control Classic conditions, respectively. Because BeatSaber blocks had a fixed duration of 150 s, these values indicate that participants required only about 8 s on average to initiate the next block. Expressed on a per-trial basis, the average trial duration was approximately 1.5 s in BeatSaber and just under 2 s in the two control conditions.

**Fig 8.**
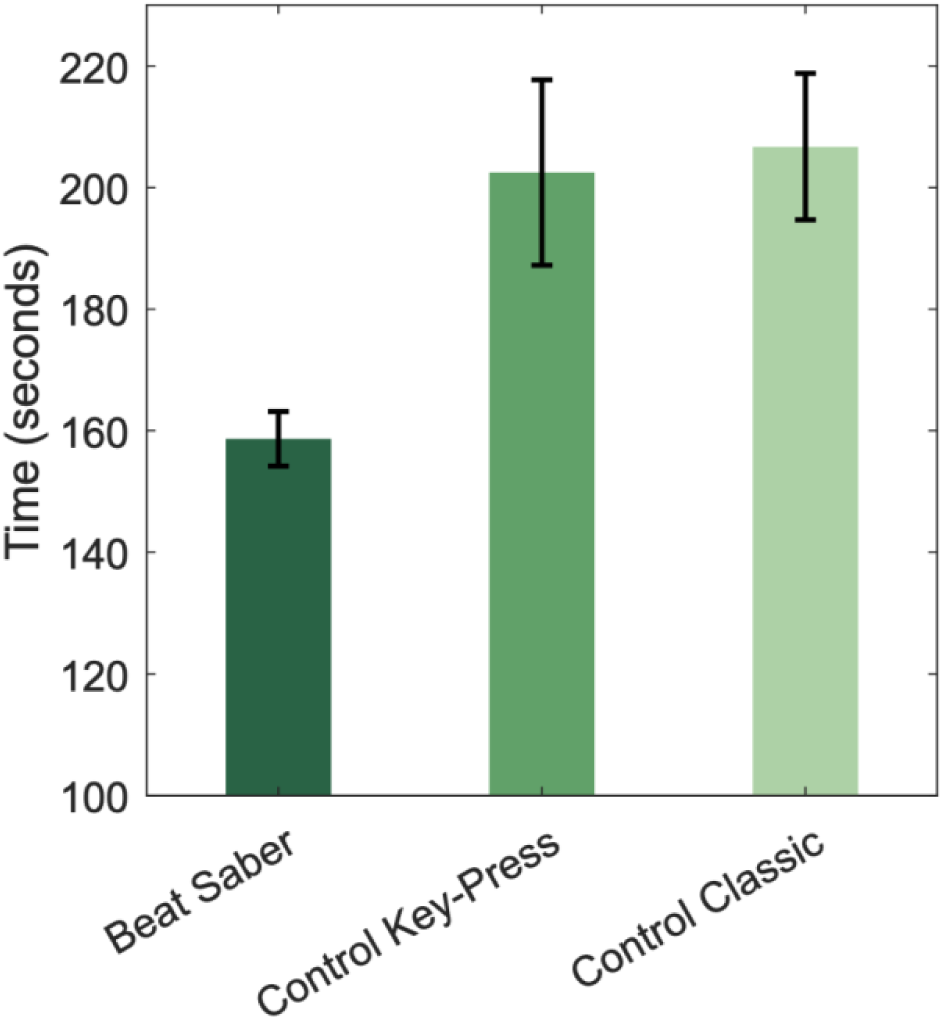
Mean time (± SEM) required to complete a block of 100 trials in the BeatSaber, Control Key-Press, and Control Classic conditions.

Despite identical stimulus presentation times across paradigms, participants completed trials more slowly in the control conditions. In these paradigms, the experiment paused after each trial and awaited a response, allowing participants to set their own pace and introducing small delays between trials. Nevertheless, the control conditions were still substantially faster than many conventional psychophysical experiments, likely because the task structure encouraged participants to maintain a continuous response rhythm. BeatSaber further increased throughput by embedding responses within a continuous stream of actions, reducing these delays and yielding the fastest overall data collection rate.

Finally, we compared Intrinsic Motivation Inventory scores between BeatSaber and the two control conditions, and User Experience Questionnaire scores between BeatSaber and Control Classic. Figure 9 shows the mean IMI scores for the ‘Interest/Enjoyment’ and ‘Pressure’ subscales, and Figure 10 shows the UEQ scores. The IMI scores reveal that participants found BeatSaber significantly more enjoyable than both Control Classic (*t*(9) = 4.185, *p* = 0.002, Cohen’s d = 1.323), and Control Key-Press (*t*(9) = 4.078, *p* = 0.002, Cohen’s d = 1.289). No difference was observed for the Pressure scale between BeatSaber and either Control Classic (*t*(9) = 2.092, *p* = 0.065, Cohen’s d = 0.661) or Control Key-Press (*t*(9) = 1.296, *p* = 0.227, Cohen’s d = 0.409). Furthermore, participants reported higher UEQ scores for BeatSaber in four of the six scales compared with Control Classic: Attractiveness (*t*(9) = 3.718, *p* = 0.004, Cohen’s d = 1.176), Novelty (t(9) = 5.378, *p* = 0.0004, Cohen’s d = 1.7007), Stimulation (*t*(9) = 4.570, *p* = 0.001, Cohen’s d = 1.445) and Efficiency (*t*(9) = 2.569, *p* = 0.0302, Cohen’s d = 0.812). We did not collect UEQ scores for the Control-Keypress condition. Overall, participants reported a more positive experience in a related questionnaire both through the scores and informal feedback when asked about their impressions.

**Fig 9.**
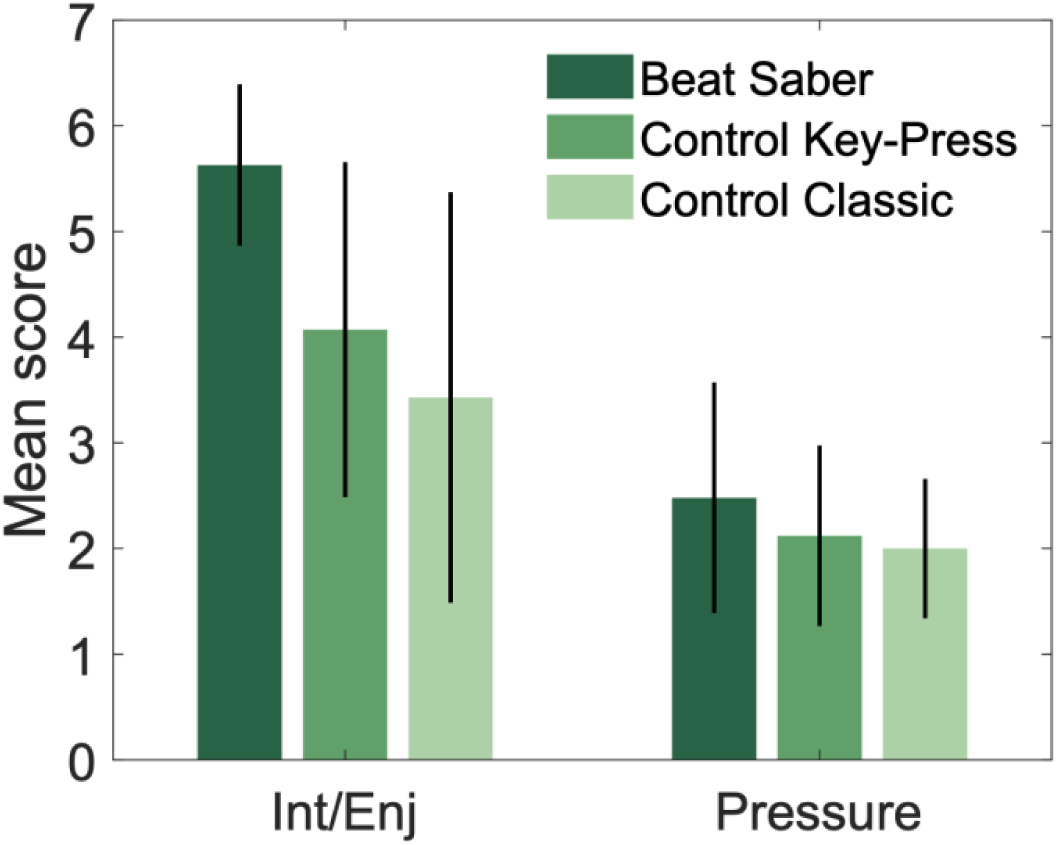
Mean scores (± SEM) for the Interest/Enjoyment and Pressure subscales of the Intrinsic Motivation Inventory (IMI) following the BeatSaber (dark green), Control Key-Press (medium green), and Control Classic (light green) tasks. Items were scored on a Likert scale.

**Fig 10:**
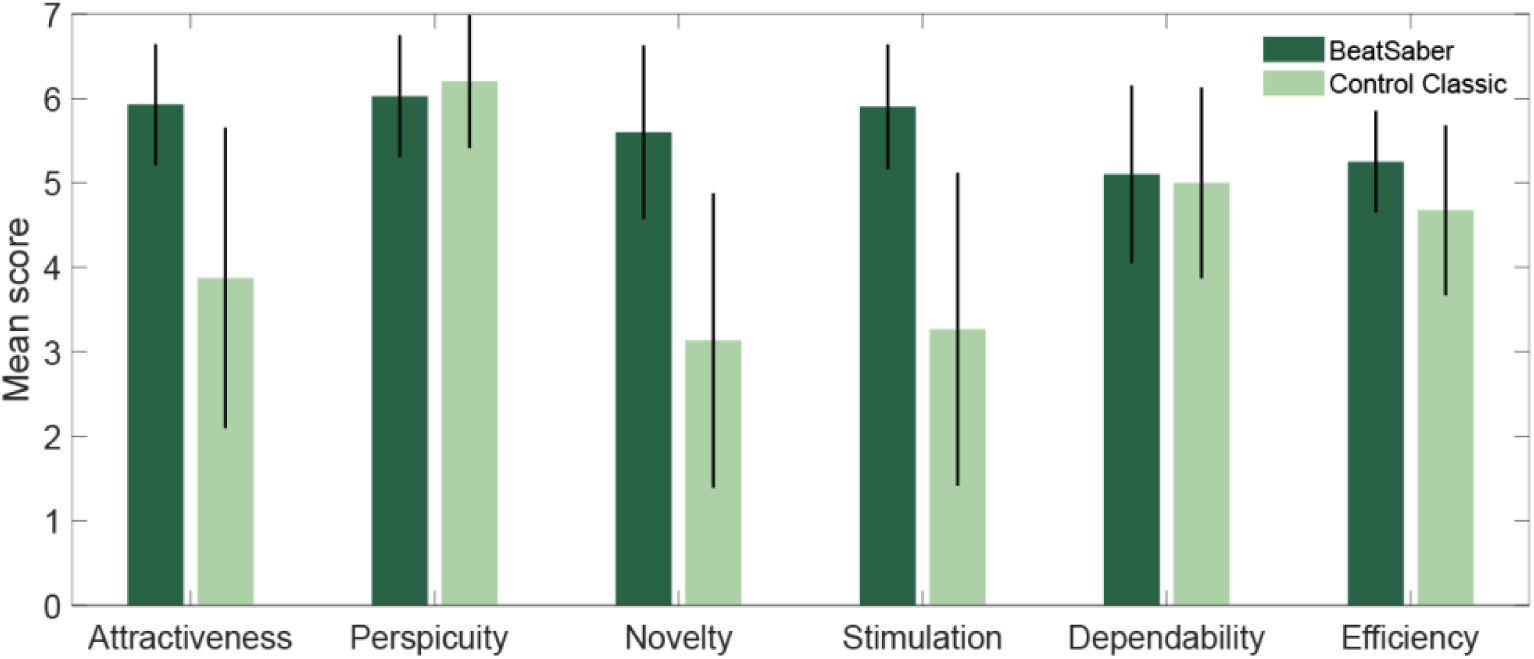
Mean scores (± SEM) for subscales of the User Experience Questionnaire (UEQ) following the BeatSaber (dark green) and Control Classic (light green) tasks. Items were scored on a Likert scale.

These results highlight two important aspects of the BeatSaber paradigm. First, psychometric performance was comparable between BeatSaber and Control Key-Press, demonstrating that an interactive, game-based environment can yield reliable threshold estimates. Second, participants consistently reported greater enjoyment and a more positive user experience than in the Control Classic condition. Together, these findings suggest that immersive, game-based psychophysics can improve participant engagement while preserving measurement quality.

## General discussion

Mapping discrimination across color space remains challenging because collecting large, high-quality psychophysical datasets is time-consuming. Most existing measurements have therefore been obtained under highly controlled laboratory conditions and remain relatively sparse. In this study, we explored whether an immersive VR paradigm could increase data throughput while preserving the quality of psychophysical measurements by embedding a standard odd-one-out 4AFC task within an engaging game environment.

Using this approach, we replicated several well-established features of color discrimination. In the DKL isoluminant plane, discrimination contours were elongated predominantly along the radial (constant-hue) direction, indicating greater sensitivity to hue than to chroma changes. Moreover, this anisotropy varied systematically across color space, being stronger for orange than for purple reference colors, consistent with previous reports (M. V. Danilova & Mollon, 2014, 2016, 2020; Hedjar et al., 2025a, 2025b; Krauskopf & Gegenfurtner, 1992).

### Reliability and Validation of the BeatSaber Paradigm

The key advantage of the BeatSaber paradigm is its high data throughput, i.e. more trials per unit time. Unlike traditional psychophysical procedures, the task proceeds continuously and does not pause while awaiting responses. Despite this rapid pace, participants maintained stable performance over extended periods, and threshold estimates were comparable to those obtained in the control conditions. BeatSaber trials lasted 1.5 s on average, compared with approximately 2 s in the two control paradigms. These values are lower than the 3 s for color-discrimination studies using conventional methods, where the program waits for a user response before continuing to the next trial (e.g., <u>Hedjar et al., 2025)</u>.

Furthermore, the present implementation was intentionally conservative. Thresholds were estimated using a method-of-constant-stimuli design with 15 repetitions at each of 10 stimulus levels. Combining the paradigm with adaptive procedures such as QUEST+ (Watson, 2017) would likely reduce the number of required trials substantially. Under such conditions, a single threshold could plausibly be estimated in roughly one minute, making dense sampling of discrimination thresholds across multiple regions of color space practically feasible.

Despite the rapid throughput and active nature of the task, the BeatSaber paradigm produced psychophysical measurements that were highly consistent with those obtained in the matched Control Key-Press condition. Because the two paradigms differed only in the response modality (controller-based slicing versus keyboard responses), this comparison provides a direct assessment of whether the motor demands of the task influence threshold estimation. Across conditions, discrimination thresholds showed close agreement, indicating that the additional motor component introduced little measurable bias. Furthermore, goodness-of-fit analyses showed that psychometric functions provided an adequate description of the data in all three paradigms, with rejection rates comparable to those typically reported in psychophysical studies (Jäkel & Wichmann, 2006). Together, these findings suggest that the BeatSaber paradigm yields reliable color-discrimination measurements despite its immersive and action-based nature.

A notable feature of the BeatSaber paradigm is that improved engagement was accompanied by stable psychophysical performance. Traditional psychophysical experiments often trade participant comfort and enjoyment for experimental control, particularly when large numbers of trials are required. By contrast, participants consistently reported that the BeatSaber task was more enjoyable than the Control Classic condition, while not experiencing greater levels of pressure or stress. The User Experience Questionnaire further indicated that participants found the task more attractive, stimulating, and novel. Importantly, these subjective benefits were not achieved at the expense of measurement quality: threshold estimates remained consistent with those obtained in the control conditions, and lapse rates were very low throughout the experiment. Together, these findings suggest that immersive, game-based paradigms may offer a practical way to increase participant motivation and sustain attention during extended psychophysical testing.

### Color Perception in Action

An immersive, gamified VR paradigm offers an opportunity to examine perceptual measurements under motor-engaged conditions. In everyday vision, perceptual judgments are rarely made in isolation but are part of ongoing action and interaction with the environment. Traditional psychophysical paradigms deliberately remove many of these factors in order to isolate specific perceptual mechanisms. While this strategy has yielded foundational insights into visual processing, it remains important to establish whether the resulting measurements generalize beyond highly constrained laboratory settings.

In the present task, participants made perceptual decisions through coordinated whole-arm movements within a dynamic and immersive environment. Despite these additional motor demands, we replicated canonical features of color discrimination, including the super-importance of hue and its systematic variation across color space. The geometric structure of color discrimination therefore appears remarkably stable across very different behavioral contexts. This robustness is noteworthy because the task transformed a perceptual judgment that is traditionally reported by a button press into a continuous, goal-directed action.

More broadly, this finding bears on longstanding discussions concerning the relationship between perception and action. Classical accounts proposed partially distinct visual pathways for perceptual judgments and visually guided actions (Goodale et al., 1994; Goodale & Milner, 1992; Milner & Goodale, 2008; Mishkin et al., 1983). At the same time, accumulating evidence indicates extensive interaction between perceptual and motor processes during natural behavior (Franz & Gegenfurtner, 2008; van Polanen & Davare, 2015). Although our experiment was not designed to adjudicate between these views, it demonstrates that one of the most robust phenomena in color vision remains largely unchanged when perceptual decisions are embedded within immersive, goal-directed action. This observation supports the broader idea that fundamental properties of color perception are preserved across a range of behavioral contexts.

### Rhythmic Interaction

So far, we have shown that the BeatSaber paradigm provides reliable psychophysical measurements while remaining intuitive, easy to learn, and highly engaging. One intriguing feature of the task is its rhythmic structure. Human perceptual and motor systems are strongly influenced by temporal regularities, and auditory rhythms are known to facilitate movement coordination and motor entrainment (Large et al., 2002; Thaut et al., 1998, 2002). In our paradigm, stimuli appeared at a fixed tempo and participants quickly synchronized their actions to this rhythm. Although we did not directly measure the contribution of rhythmic entrainment, it may help explain why observers were able to perform precise perceptual judgments using full-arm movements without any apparent loss of psychophysical sensitivity.

This observation raises an interesting possibility for future research. Rather than treating motor responses as an unavoidable component of perceptual experiments, rhythmic and game-based paradigms may actively support rapid and consistent responding. Such approaches could provide a natural framework for implementing speeded perceptual decisions while maintaining participant engagement over extended testing sessions. Combined with the increased throughput demonstrated here, this may make immersive and gamified paradigms particularly valuable for collecting the large datasets required for modern psychophysics.

### Limitations and Practical Considerations

There are, of course, limitations to immersive and action-based psychophysical tasks. The BeatSaber paradigm requires a basic level of motor coordination and a short period of practice, and it may therefore be less suitable for some participant populations than traditional button-press paradigms. Whereas conventional psychophysical experiments are often limited by attentional lapses and fatigue, active tasks may be more susceptible to motor errors that can be misinterpreted as perceptual mistakes. Careful task design and sufficient practice are therefore important for balancing these trade-offs.

Importantly, the core principles of the paradigm are not restricted to virtual reality. Similar interactive tasks could be implemented on conventional displays, potentially combining some of the engagement benefits of gamified psychophysics with the experimental control and accessibility of traditional laboratory setups. Exploring such approaches may be particularly useful in contexts where VR equipment is unavailable or impractical.

A second set of limitations concerns the display technology itself. As with any visual experiment, the optical and photometric properties of the display place constraints on stimulus generation and interpretation. Current VR headsets generally do not yet match the calibration accuracy, spatial resolution, or optical quality of the best laboratory displays used in vision research. Nevertheless, display technology continues to improve rapidly, and our results add to a growing body of evidence demonstrating that modern VR systems can support rigorous studies of color perception (Barrancas et al., 2023; Gil Rodríguez et al., 2022; Rodríguez et al., 2024).

### Towards Active Psychophysics

This work is part of an emerging framework in perceptual research that aims to study sensory and decision-making processes through more active behavior. Continuous psychophysics (Barnett et al., 2025; Bonnen et al., 2015, 2017; Huk et al., 2018; Knöll et al., 2018; Straub & Rothkopf, 2022) has been an influential approach to this. By moving past the rigid structure of independent trials and instead eliciting continuous behavioural adjustments to a dynamic stimulus, it allows researchers to model perception as an ongoing process guiding action. Virtual Reality provides complementary advantages by enabling immersive, interactive environments while preserving rigorous control over stimulus presentation (de Gelder et al., 2018; Tachibana & Matsumiya, 2022; Wilson & Soranzo, 2015). Studying color perception with a ‘BeatSaber’ like game was first pioneered by Aizenman et al. (2026) to measure color category boundaries. Our work builds on this foundation by demonstrating that such an approach can be extended to the quantitative measurement of color discrimination thresholds, while formally validating both data quality and participant engagement.

## Conclusion

By enabling faster and more engaging data collection without compromising measurement quality, immersive paradigms offer a practical route toward constructing more comprehensive empirical maps of perceptual color space. Approaches of this kind may help overcome the sparse sampling that has historically constrained the field, bringing large-scale metric characterization of color space within practical reach. As display technologies continue to improve, immersive and action-based psychophysical methods may help bridge the gap between controlled laboratory experiments and natural behavior, opening new opportunities to study color perception not only in the laboratory, but in the context of the actions for which vision evolved.

## Acknowledgments

We would like to thank Laysa Hedjar for discussion and advice. We would also like to thank Tugce Kocancioglu for administrative support with the project.

## Data availability

All data will be deposited on Zenodo upon acceptance of the manuscript.

